# Evidence of enzyme-level thermal constraint on biological nitrogen fixation rates across systems and scales

**DOI:** 10.1101/2025.06.17.660177

**Authors:** Kaleigh E Davis, Margaret A Slein, Mary I O’Connor

## Abstract

Biological nitrogen fixation (BNF) provides half of global new nitrogen annually and plays an important role in biodiversity patterns and global biological carbon uptake processes. BNF rates accelerate with warming, with known implications for ecological functioning, yet the strength of this temperature sensitivity and its context dependence are not well understood. Here we synthesized 70 controlled experimental tests of the acute temperature dependence of nitrogen fixation rates and found that BNF rates accelerate with temperature in a consistent way across levels of biological organization from enzyme to community, mirroring scaling of metabolic temperature dependence across levels of organization for respiration, photosynthesis and methane production. This BNF temperature dependence is also remarkably consistent across biological systems. This analysis shows multiple lines of evidence for general, scalable effects of temperature on nitrogen fixation rates, in line with previous suggestions of a strong enzyme-level constraint. This widespread pattern may be important for understanding effects of warming on coupled carbon-nitrogen systems with ongoing global change.

## Introduction

Despite its vast abundance on the planet, nitrogen is one of the most common limiting nutrients in nature (Elser *et al*. 2007; Falkowski 1997; LeBauer & Treseder 2008). Biological nitrogen fixation (BNF) converts inert dinitrogen (N_2_) to bioavailable ammonium (NH_4_^+^) and is a critical metabolic link between the vast stores of atmospheric nitrogen and the biosphere. Nitrogen from BNF plays an important role in biogeographic diversity patterns and global biological carbon uptake processes by facilitating primary production in low and variable nitrogen environments (Brookshire *et al*. 2019; Houlton *et al*. 2008; Lu *et al*. 2019; Steidinger *et al*. 2019). Indeed, the relationship between BNF and oxygenic photosynthesis is nearly as old as photosynthesis itself (Knoll 2008; Latysheva *et al*. 2012), and close association with nitrogen fixation is common across the tree of photosynthetic life, including cyanobacteria, protists, lichens, and lower and higher plants (Caputo *et al*. 2019; Postgate 1982). As climate warming accelerates metabolic demand across all systems, it is not clear if there are general patterns for how nitrogen production from BNF will respond to warming. A deeper understanding of these effects is needed to understand changes in ecological functioning with warming (Thornton *et al*. 2007).

It has long been understood that, because breaking the triple bond in a dinitrogen molecule is an energy intensive process, nitrogen fixation rates are highly temperature sensitive (Burns 1969; Ceuterick *et al*. 1978). The strong temperature sensitivity of BNF has been invoked to suggest that BNF is a more viable nitrogen acquisition strategy at higher temperatures (Houlton *et al*. 2008), and in turn to explain ecological phenomena including spatio-temporal variation in the occurrence of cyanobacteria blooms (Jöhnk *et al*. 2008) and the geographic distribution of nitrogen fixation in marine pelagic (Breitbarth *et al*. 2007; Staal *et al*. 2003; Ward *et al*. 2013) and terrestrial systems (Houlton *et al*. 2008; Steidinger *et al*. 2019; Wang & Houlton 2009). These explanations have proven useful for describing ecological processes in diverse systems, but without a common mechanistic understanding for how warming affects nitrogen fixation across scales, the field is left with disparate predictions for how the productivity of nitrogen-fixing systems, and larger scale couple N- and C-budgets, will change with warming (Bytnerowicz *et al*. 2022; Houlton *et al*. 2008; Wang & Houlton 2009; Williamson *et al*. 2016). Resolving how fundamental relationships between BNF and temperature link processes across ecological contexts is thus a key outstanding question in climate science (Deutsch *et al*. 2024).

Across its broad ecological and taxonomic domain, BNF is mediated by a single enzyme, nitrogenase, with minimal structural and biochemical variation among taxa (Emerich & Burris 1978; Harris *et al*. 2019; Raymond *et al*. 2004; Zehr *et al*. 2003). From this, a mechanistic hypothesis is that the effect of temperature on the kinetics of nitrogenase activity also varies minimally across organisms employing this common biochemistry (Deutsch *et al*. 2024; Gillooly *et al*. 2006). Further, if enzyme-level thermal responses constrain thermal responses at higher level of biological organization, as suggested by metabolic scaling theory (cf. Brown *et al*. 2004) and ecological constraint hypotheses (Deutsch *et al*. 2024; Rezende & Bozinovic 2019), we could expect to see similar temperature effects on BNF rates at organism, population, and ecosystem levels. Indeed, general effects of temperature on metabolic rate across biological systems and levels of organization have been observed for other highly conserved metabolic pathways, including aerobic respiration (Gillooly *et al*. 2001; Smith *et al*. 2021), photosynthesis (Lopez-Urrutia *et al*. 2006; Yvon-Durocher *et al*. 2010), and methanogenesis (Yvon-Durocher *et al*. 2014). Recently, this enzyme-constrained scaling relationship has been invoked to explain restricted variation in thermal optima for BNF across ecosystem types (Deutsch *et al*. 2024). However, many systems operate well below their thermal optima, in the region of the thermal niche where the rate of acceleration with warming determines ecological outcomes (Bernhardt *et al*. 2018; Vasseur *et al*. 2014) (Fig 1A). To understand how BNF-supplied nitrogen will change with warming, a key question is whether warming-induced acceleration of BNF rates at temperatures cooler than systems’ thermal optima show evidence of enzyme-level constraint, and therefore cross-system predictability.

**Figure 1.**
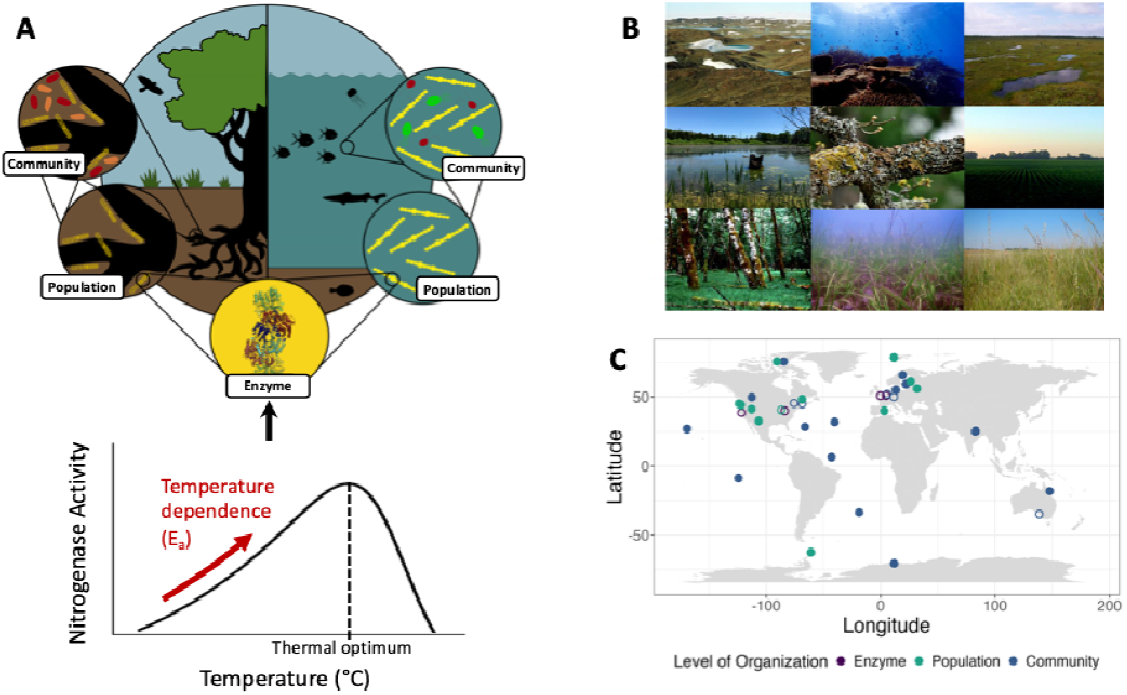
Nitrogen fixation across diverse ecological contexts. A) Diagram showing the ubiquitous nitrogen fixing enzyme, nitrogenase, and its thermal response. This enzyme-level thermal response is hypothesized to underpin nitrogen-fixation thermal responses at higher levels of organization, shown here in circles, and across ecosystems (two domains shown as two halves of the largest circle). Our analysis handles temperature dependence, as it is characterized in the increasing portion of the thermal response. B) A subset of the diverse ecosystems represented by our nitrogen fixation temperature-response dataset occur, including an Antarctic ice-free plateau, coral reef, sphagnum bog, freshwater pond, lichen, soybean crop field, alder grove, seagrass meadow, and prairie grassland. C) Geographical distribution of diazotrophs represented in this study. Filled circles display the location of *in situ* experiments or the location of the original source site if the study used isolated strains for laboratory experiments. Open circles display the location of the corresponding author’s research institution if no information on source location was not available. Image credits: Nitrogenase enzyme structure - RSPDB M1N1; antarctic plateau - Dr. Amit Dharwadkar; sphagnum bog - Wikimedia commons; soybean field - Wikimedia commons; freshwater pond – Dr. Jason Laurich; lichen - Andrew Simon; prairie grassland - Wikimedia commons.

The metabolic scaling approach provides an efficient first order assessment of whether simple models of temperature effects on enzyme (here, nitrogenase) activity can describe broad-scale temperature effects on metabolic (here, BNF) rates. This model would predict that nitrogenase activity responds similarly to warming across systems, regardless of taxonomic or ecological context, and that this enzyme-level thermal response scales to higher levels of biological organization (cf. Brown *et al*. 2004; Enquist *et al*. 2003; Gillooly *et al*. 2001; Houlton *et al*. 2008). However, nitrogen fixation occurs across phylogenetically distinct taxa in very different biological contexts, including terrestrial and aquatic habitats, free-living and symbiotic systems, and in close association with and independent from photosynthetic systems. These differences may impose structural or genetic constraints on nitrogen-fixing organisms and may fundamentally alter the rates of energy supply and demand to them, leading to thermal responses of BNF in different systems or at higher levels of biological organization that may diverge from enzyme-level patterns rather than scale directly (Deutsch *et al*. 2024; Yvon-Durocher *et al*. 2014) (Figure 1). If these ecological and evolutionary differences among nitrogen-fixing systems alter the temperature-dependence of BNF then models more complex than a simple scaling of enzyme-thermal-kinetics may be needed to understand changes in BNF rates with warming.

Here, we synthesized the most comprehensive dataset to date of observed BNF temperature responses to assess patterns in the temperature dependence of BNF across ecosystems and levels of biological organization. We draw from metabolic scaling theory to formulate and test three hypotheses. First, we tested a **general temperature dependence hypothesis** that the temperature response of nitrogen fixation is similar across ecological systems, because it is underpinned by a similar kinetic response of the enzyme nitrogenase in each of these systems. We tested this against an alternative **systematic temperature dependence hypothesis**, that the temperature dependence of BNF varies with ecological or evolutionary context. Specifically, we tested for systematic variation in BNF temperature dependence with: nitrogen fixer (hereafter “diazotroph”) taxonomy, habitat type, photosynthetic association, and thermal acclimation history. We tested these hypotheses at each level of organization independently, which allowed us to identify potential drivers of deviation from enzyme-level constraints as systems become more complex. Finally, we tested a **metabolic scaling hypothesis**, that a common enzyme-level BNF temperature-dependence constrains BNF temperature responses at population and community levels. This hypothesis predicts that temperature dependencies at these higher levels of organization will scale with the enzyme-level response. In testing these three hypotheses, we determine whether simple scaling relationships can be used to describe temperature-driven change in nitrogen fixing systems at non-stressful temperatures, and whether the domain of metabolic scaling theory can be extended to include a fourth metabolic process.

## Methods

We synthesized a dataset of 70 unique BNF thermal-responses from the published literature and estimated the effect of temperature on BNF-rate independently at enzyme, population, and community levels by using linear mixed effects models to fit the linearized Boltzmann-Arrhenius function to datasets at each level of organization. We then compared the general and variable temperature dependence hypotheses using AICc-based model selection (Yvon-Durocher *et al*. 2014). Thus, for the purposes of this study, general versus variable temperature dependence are considered mutually exclusive, where the temperature dependence of BNF at each level of organization is either consistent across studies or variable with some ecological or evolutionary factor(s) identified in the dataset (Table S1). We tested our metabolic scaling hypothesis using two analyses in concert; first, we compared model estimates from the enzyme-, population-, and community-level analyses, and second, we employed a cross-level, whole-dataset model comparison approach.

### Data Synthesis

We tested the three hypotheses using data from published studies that presented BNF rate measurements over an experimental temperature gradient (hereafter, “thermal responses”). We identified papers in Web of Science using the search terms “nitrogen fix*” and “temperature.” We screened 5143 papers first by title, then by abstract using the revtools package in R (Westgate 2019 R version 4.0.2). We conducted a full text review for 311 papers that indicated in the abstract the reporting of nitrogen fixation rate measurements across multiple temperatures. We applied three criteria to determine whether data were appropriate for inclusion in our analysis: 1) all BNF rate measurements were mass-corrected (e.g., units N fixed per unit mass per unit time), 2) BNF rate measurements were made over a minimum of four temperatures below the thermal optimum (see Figure 1A) and 3) temperature measurements were instantaneous (i.e., not averaged over time or space). In applying these criteria, we excluded 284 papers that could not be used for our analysis, resulting in a full dataset consisting of 70 unique BNF thermal responses from 27 studies (Table 1).

**Table 1.** Distribution of temperature-responses across ecological and evolutionary factors tested in our main analysis. The sample size given in next to each level of organization refers to the number of thermal responses in the dataset.

| Level of organization (n) | Factor<sup>†</sup> | Level | No. of temperature-responses |
| --- | --- | --- | --- |
| Enzyme (4) | Nitrogenase source species | Azotobacter vinelandii | 2 |
|  |  | Klebsiella pneumoniae | 2 |
| Population* (39) | Habitat | Aquatic | 18 |
|  |  | Terrestrial | 21 |
|  | Diazotroph Order | Nostocales | 18 |
|  |  | Oscillatoriales | 7 |
|  |  | Hyphomicrobiales | 7 |
|  |  | Chroococcales | 3 |
|  |  | Actinomycetales | 2 |
|  |  | Rhodospirillales | 1 |
|  |  | Pseudomonadales | 1 |
|  | Thermal acclimation history | Acclimated | 14 |
|  |  | Not Acclimated | 25 |
| Community* (27) | Photosynthetic association | Yes | 10 |
|  |  | No | 17 |

<sup>†</sup> See Table S1 for definitions of each factor and level
\* See Figures S4 and S5 for overlap between these factors at the population and community levels, respectively

Diazotroph biomass was quantified and reported, usually as a part of the composite mass-corrected rate variable, by the original study authors. We requested raw data from authors and extracted data from papers using WebPlotDigitizer (Rohatgi 2022) when author-sourced data were unavailable. In the final dataset, all BNF rate measurements were conducted using the acetylene reduction assay method (See Shao *et al*. 2023 for comparison with direct tracing methods). We categorized each BNF temperature response by the level of biological organization (i.e., enzyme, population, or community) at which data were collected, to test our scaling hypothesis. Here, population and community are used in the ecological sense, to refer to coexisting organisms of the same species, and coexisting organisms of differing species, respectively. Enzyme-level studies were identified by the isolation of nitrogenase enzyme for empirical estimation of BNF rate. Studies were categorized as population-level if a single nitrogen-fixing taxon was identified in the study and categorized. Studies were categorized as community-level if the diazotrophs were described in the research article as multi-species or if nitrogen fixation was measured on an unmanipulated sample of a natural substrate, such as a soil core or segment of a log (Table S1).

### Model Fitting

To test the general temperature dependence hypothesis and alternative variable temperature dependence hypothesis, we fit thermal response models to each set of BNF rate measurements across a temperature gradient (hereafter BNF temperature-response), at each level of biological organization. Metabolic temperature responses are generally non-linear and unimodal (Angilletta Jr 2009; Arroyo *et al*. 2022; Figure 1A). Fitting unimodal models to accurately describe both the increase in rates at low temperatures and the decline in rates at high temperatures requires high resolution data at temperatures above and below the thermal optimum (Michaletz & Garen 2024). From our full dataset of 70 thermal responses, only 41 (59%) thermal responses contained at least one data point at a temperature beyond an apparent thermal optimum, and fewer than five studies contained higher resolution data past an apparent thermal optimum. Thus, fitting a unimodal model was inappropriate (Michaletz & Garen 2024). Instead, we truncated thermal response data to include data in the sub-optimal thermal range only, and fit thermal responses using an exponential model (Michaletz & Garen 2024), which characterizes the accelerating effect of temperature on biological processes up to temperatures approaching the thermal optimum (Figure 1A). For our analysis, we used a linearized form of the widely used Arrhenius model:

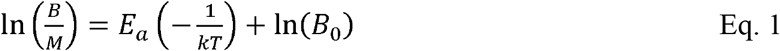

where the biomass-corrected nitrogen fixation rate, 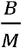 (e.g. g N fixed / hr /g diazotroph biomass), is related linearly to inverse temperature, 1/(*kT*) (K), via Boltzmann’s constant, *k* (eV/K), the temperature dependence parameter, *E*_*a*_ (eV), and the model intercept, *B*_0_(units N fixed per unit mass per unit time). The parameter, *E*_*a*_, represents the slope of the log rate-temperature relationship and is the focal parameter of our analysis.

We estimated the value of *E*_*a*_ for each BNF temperature-response using linear mixed effects models that took the following form:

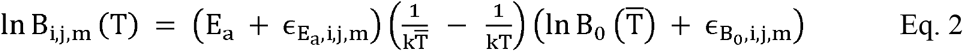

where *B*_*i,j,m*_ is the natural log of biological nitrogen fixation rate at temperature, *T*(K), for some observation, *i*, in each experimental unit, *j*, in study, *m*. Each experimental unit corresponds to a set of BNF rate measurements made on replicate enzymes, populations or communities for which all conditions were held constant except for temperature, referred to as a ‘temperature response’ in the main text. 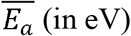 captures the temperature dependence estimated over all experimental units within the level of organization being analyzed. We centered temperature data using the mean temperature across all observations at each level of organization, 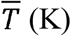 at each level of organization. ln 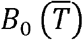 represents an average of BNF rate observations at mean temperature, 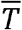. The model allowed for variation in *E*_*a*_ among experimental units and among studies, and modeled this variation using a random effect 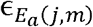 with a normal distribution with a mean 0 centered on 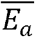 (following O’Connor *et al*. 2007; Yvon-Durocher *et al*. 2014). We also included variation among experimental units and studies in the intercept term ln 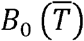, which reflects variation in system-level BNF rates, experimental conditions, and units in which BNF rates were reported. We did not standardize rate units across studies as comparisons of model intercepts were not relevant to our hypotheses.

To test the general temperature dependence hypothesis against the variable temperature dependence hypotheses, we compared the general temperature dependence model given by Eq. 2 to expanded models (e.g. Equation S1) with temperature interaction terms for ecological and evolutionary factors expressed as fixed effects (Table S1). The factors we tested in the main analysis were: diazotroph species at the enzyme-level; diazotroph Order, habitat, thermal acclimation history at the population level; and association with photosynthetic organisms at the community level (Table 1). While we expected that traits such as oxygen accommodation strategy (heterocysts, temporal C/N separation, or none) or symbiosis status (symbiotic or free-living) might also be sources of systematic variation, all variation among taxa in symbiosis vs free-living life histories (and associated variation in ability to directly estimate diazotroph biomass) and in oxygen accommodation strategy was captured by Order, indicating complete phylogenetic co-variation among these factors in our dataset (Figure S1). At the community level, unbalanced data availability across habitat types precluded us from testing habitat as a source of variation (Figure S2). All models included random effects of BNF temperature response nested within study on model slope and intercept (e.g. Equation S1) and were fit using the nlme package in R (Pinheiro *et al*. 2023).

### Model selection and inference

To determine whether a general temperature dependence or variable temperature dependence better described BNF temperature responses, we used small-sample-corrected AIC (AICc)-based model comparison with a threshold ΔAICc value of 2 (Burnham & Anderson 2004) executed in the R package MuMIn (Bartoń 2009). We used the temperature terms retained in the best model to make inferences about the two alternative hypotheses: if no temperature interaction terms were included in the best model, we inferred support for a general temperature dependence; if at least one temperature interaction term was included in the best model, we rejected the general temperature dependence hypothesis (Garzke *et al*. 2019; Yvon-Durocher *et al*. 2012). If the best model contained a temperature interaction term(s), we compared that model against a model that allowed only the intercept to vary with factor(s) in the interaction term(s). Because model intercepts were not relevant to our hypotheses, if the best model included an effect of a factor on model intercept, but not on model slope (i.e. a temperature by factor interaction), we inferred evidence for the general temperature dependence hypothesis. We first fit models using maximum likelihood estimation to facilitate comparison of models with different fixed effects structures, then refit the best model using restricted maximum likelihood estimation to estimate model coefficients (Zuur *et al*. 2009). We conducted this model comparison analysis separately at each level of organization.

To test the metabolic scaling hypothesis, we took a two-pronged approach: first, we compared the 95% confidence intervals around model-estimated temperature dependencies at population and ecosystem levels with that at the enzyme level; second, we employed a cross-level model comparison approach to test whether a single temperature dependence term (*E*_*a*_) across the whole dataset described the temperature dependence of N-fixation within a level of organization better than separate temperature dependence terms at each level. For each test, respectively, we inferred evidence consistent with metabolic scaling if confidence intervals included the enzyme-level estimate, and if the best model included only a temperature term and no temperature by level of organization interaction. All analyses were conducted using R version 4.3.2.

## Results

An analysis of BNF temperature responses from 70 controlled experiments within 27 studies revealed a consistent suboptimal BNF temperature dependence of 1.03 ± 0.13 eV from enzyme to community levels (Figure 2, Figure 3, Figure S3). We found stronger support for a general temperature dependence across systems than for variable temperature dependencies at population and community levels, and mixed support at the enzyme level (Table 2). The enzyme-level BNF temperature dependence estimate was included within 95% confidence intervals for all higher-level temperature dependence estimates, in support of the metabolic scaling hypothesis (Figure 3). A second full-dataset analysis provided a second line of evidence supporting the metabolic scaling hypothesis. We compared temperature dependence estimates for linearized exponential and nonlinear (quadratic) models, which account for decelerating rates of BNF near an optimal temperature (Figure 1A; Pawar *et al*. 2016 Eq. 10), and found that they produced nearly identical temperature dependence estimates (Figure S4). All subsequent results employ the linearized Arrhenius expression of temperature dependence, as this model has wide use in metabolic scaling theory and the temperature-dependence term, *E*_*a*_, is anchored in first principles of temperature effects on enzyme kinetics.

**Table 2.** Results of model selection to determine systematic covariates of temperature effects on BNF rate at each level of organization. Best models at each level of organization are displayed in bolded text and models are presented within each level in order of AICc value. Temperature is abbreviated to “temp” and thermal acclimation history is abbreviated to “therm acclim hist.” Order refers to the taxonomic order of the diazotroph, as identified in the source study. Full descriptions of the definitions and the sorting criteria for each of the fixed effects are presented in Table S1. All models include a random slope and intercept for experimental unit, and in the population- and community-level datasets, random effects of experimental unit are nested within study.

| Level of organization | Fixed effects structure | df | logLik | AICc | ΔAICc | Weight |
| --- | --- | --- | --- | --- | --- | --- |
| **Enzyme** | **Temp** | **6** | **-35.5** | **86.3** | **0.00** | **0.58** |
|  | Temp * species | 8 | -32.8 | 87.7 | 1.39 | 0.29 |
|  | Temp + species | 7 | -35.4 | 89.2 | 2.92 | 0.13 |
| **Population** | **Temp + order** | **15** | **-256.0** | **544.8** | **0.00** | **0.83** |
|  | Temp * therm acclim hist + order | 17 | -255.7 | 549.0 | 4.18 | 0.10 |
|  | Temp * order + temp * therm acclim hist | 23 | -248.9 | 550.6 | 5.78 | 0.05 |
|  | Temp * order | 21 | -253.2 | 554.0 | 9.13 | 0.01 |
|  | Temp * habitat + temp * order + temp * therm acclim hist | 25 | -248.0 | 554.1 | 9.31 | 0.01 |
|  | Temp * order + therm acclim hist | 22 | -253.0 | 556.2 | 11.3 | 0.00 |
|  | Temp * habitat + temp * order | 23 | -252.6 | 558.0 | 13.2 | 0.00 |
|  | Temp + therm acclim hist | 10 | -271.8 | 564.8 | 19.9 | 0.00 |
|  | Temp | 9 | -273.5 | 565.9 | 21.1 | 0.00 |
|  | Temp * therm acclim hist | 11 | -271.5 | 566.5 | 21.7 | 0.00 |
|  | Temp * habitat | 11 | -273.1 | 569.7 | 24.8 | 0.00 |
|  | Temp * habitat + temp * therm acclim hist | 13 | -271.0 | 570.0 | 25.2 | 0.00 |
| **Community** | **Temp** | **9** | **-266.5** | **551.9** | **0.00** | **0.88** |
|  | Temp * photosynthetic association | 11 | -266.2 | 556.0 | 4.00 | 0.12 |

**Figure 2.**
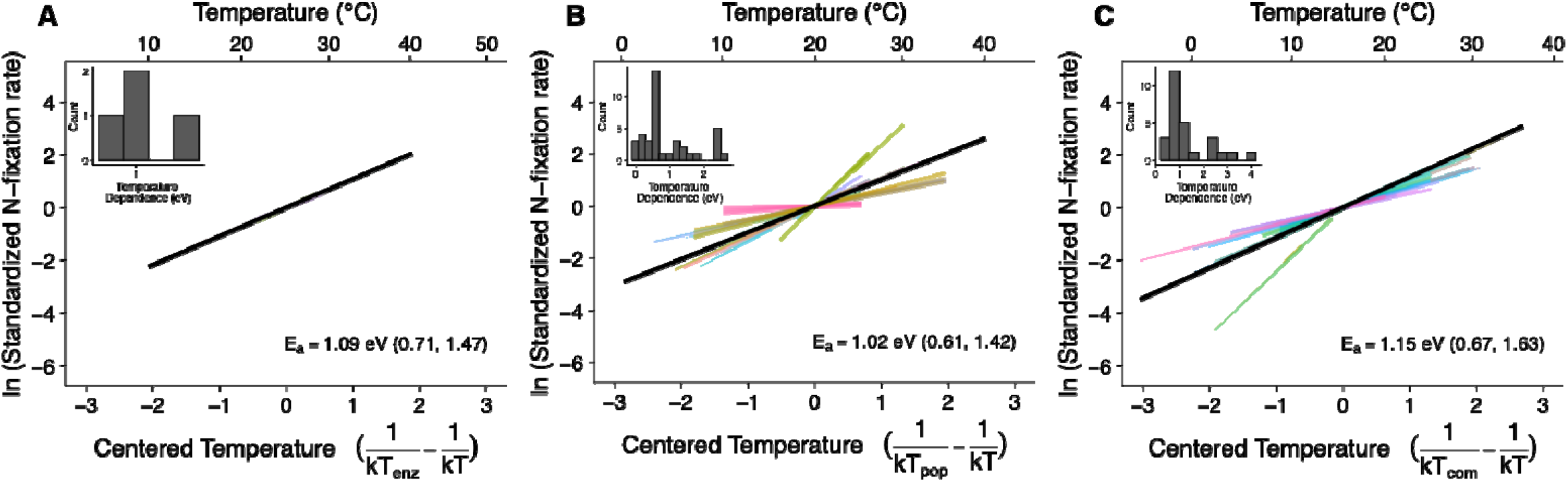
General temperature dependence of BNF across three levels of organization, A) enzymes, B) populations, and C) communities, based on hierarchical mixed effects model analysis. Model-predicted BNF rate-temperature relationships (black lines) and random slopes fit to each thermal response (colored lines). Lines color corresponds to study. Note that in the panel A, two of the four thermal response lines are hidden by the model-predicted enzyme-level line. Nitrogen fixation rates are standardized to the predicted rate at the mean temperature for the level of organization to facilitate visual slope comparison. Insets show the distribution of these model fits including random effects at each level of organization. Model estimates are displayed over raw data in Figure S3.

**Figure 3.**
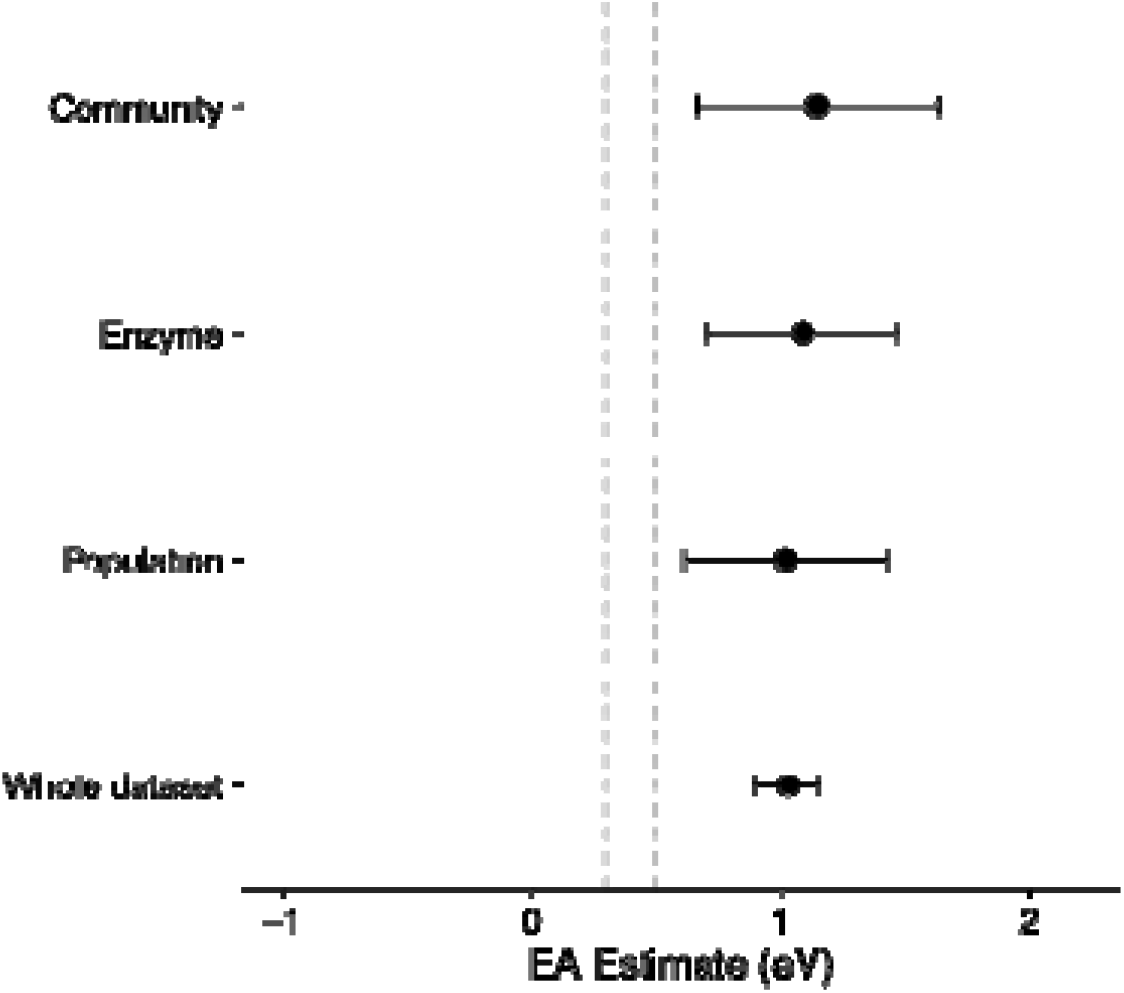
Metabolic scaling coefficients for BNF temperature dependence. Temperature dependence estimates and 95% confidence intervals are displayed from the best model at each level of organization and across the whole dataset. The grey shaded area displays the range of temperature dependence estimates commonly used for photosynthesis, 0.3-0.5 eV.

Across the enzyme-level dataset (n = 4 temperature responses), the estimated BNF temperature dependence was 1.09 eV, 95% CI: 0.71 to 1.47 eV (Figure 2A). Model comparison between a general temperature dependence model and a variable temperature dependence model revealed equivocal evidence for a single temperature dependence across these studies: the best model included only a main effect of temperature on BNF rate, but the second-best model (ΔAICc = 1.39) included variation in the temperature effect with nitrogenase source-species (Table 2). In the variable temperature dependence model, there was a significant difference in the predicted BNF temperature dependence between the two nitrogenase source species represented in the dataset: nitrogenase isolated from *A. vinelandii* (n = 2) had an *E*_*a*_ of 1.32 eV, 95% CI: 0.95 to 1.69 eV and nitrogenase isolated from *K. pneumoniae* (n = 2) has an *E*_*a*_ of 0.75, 95% CI: 0.21 to 1.29 eV (Figure S5).

For mono-specific populations of diazotrophs (n = 39 temperature-responses), our analysis revealed a BNF temperature dependence of 1.02 eV with 95% CI: 0.61 to 1.42 eV, similar to that of the nitrogenase enzyme (Figure 2B). The general temperature dependence model outperformed models that allowed temperature main effects to vary systematically with habitat, diazotroph order, or thermal acclimation history (Table 2). For comparisons of within-community temperature dependence for multispecies diazotroph assemblages, analysis (n= 27 temperature-responses) revealed a BNF temperature dependence of 1.15 eV with 95% CI: 0.67 to 1.63 eV, similar to those at the population- and enzyme-levels (Figure 2C). Here, model selection indicated stronger support for the general temperature dependence model than for a model where main effects of temperature varied with photosynthetic association (Table 2). Exceptionally strong temperature responses in population- and community-level datasets (6 in each dataset; Figure 2B and C insets) are likely attributable to study-level differences as indicated by the random effect, because the biological systems (particular species or communities) that comprise these extreme rates were represented elsewhere in the data by less extreme estimates published in different studies. Further, all extreme temperature sensitivity values in the population-level dataset were reported in a single study, and most extreme values in the community-level dataset are represented by a different single study (Figure 2).

Comparing confidence intervals on estimates from the analysis at each level of organization (enzyme, population, community) (Figure 3) suggests that BNF-temperature responses are similar across scales. Additionally, a cross-level model comparison analysis revealed that a model with a single temperature coefficient across all three levels of organization outperformed a model that fit separate slopes at each level (Table S2), and produced a BNF temperature dependence estimate of 1.03 ± 0.13 eV. Together, these two lines of evidence provide strong support for the metabolic scaling hypothesis.

## Discussion

Ecological models using metabolic scaling theory have provided simple, parsimonious explanations for several macroecological patterns from biogeographic variation in ocean carbon metabolism (Lopez-Urrutia *et al*. 2006) to biodiversity (Stegen *et al*. 2009) to methane production (Yvon-Durocher *et al*. 2014). While several studies have noted the universality of the nitrogenase enzyme in nitrogen-fixing systems (Harris *et al*. 2019; Raymond *et al*. 2004; Zehr *et al*. 2003), our analyses have provided the first strong evidence that the rate of new nitrogen production through biological nitrogen fixation (BNF) responds to temperature consistently across systems and scales. We estimated an enzyme-level BNF temperature dependence of 1.09 eV in the suboptimal temperature range that then emerged at the population and community levels, consistent with the hypothesis that the temperature dependence of the enzyme-level metabolic system scales to higher levels of organization. This BNF temperature dependence estimate is consistent with previous reports of ecosystem-level terrestrial nitrogen fixation (approximately 1.07 eV; Houlton *et al*. 2008) and substantially lower than a widely used enzyme-level estimate (2.15 eV; Ceuterick *et al*. 1978). Additionally, general temperature dependencies emerged across biological systems at the population and community levels. Together, these results provide evidence for a strong and consistent temperature dependence of nitrogen fixation across ecosystems and levels of biological organization, suggesting BNF has generalized thermal behavior constrained by enzyme-level kinetics.

We tested for a common enzyme-level temperature dependence of BNF to understand whether a thermal constraint on metabolic rate could be observed at population and community levels (Brown *et al*. 2004; Gillooly *et al*. 2001), and to contribute a rare, if small, data synthesis at this basal level of organization (but see Gillooly *et al*. 2006). While an enzyme with a consistent structure and biochemistry should theoretically have a characteristic temperature dependence, we found weak evidence that the temperature dependence of nitrogenase activity may vary depending on source species, *Azotobacter vinelandii* and *Klebsiella pneumoniae* (Table 2). This difference is surprising, because, while these two species can utilize slightly different variants of nitrogenase (Cabello *et al*. 2009), there is evidence that these nitrogenases are nearly perfectly cross-compatible (Emerich & Burris 1978; Thorneley *et al*. 1975). With only two taxa and four datasets, we are limited in the capacity to extend this inference to broader groups; a comprehensive treatment of enzyme-level temperature effects requires data from across all nitrogenase groups (Koirala & Brözel 2021; Zehr *et al*. 2003). However, this synthesis is similar in scope to other enzyme-level syntheses(e.g., Gillooly *et al*. 2006) and our findings provide a more informative prediction for enzyme-level temperature dependence than a single estimate, as is often used. Further, within this study these data serve as a useful basis for comparison for the more abundant BNF-temperature rate data at higher levels of organization.

From possible enzyme-level differences in BNF temperature response across species, we might expect to observe systematic effects of species identity on temperature dependence or much wider confidence intervals around the estimate in the population and community level datasets, which contain many taxa across diverse Orders (Table S1). Instead, we found that BNF temperature dependencies at higher levels of organization showed similar temperature dependence to the mean enzyme-level temperature dependence. Further, in our population-level analysis, (N = 39 responses) BNF temperature dependence did not vary with diazotroph taxonomy, which spanned seven Orders. This suggests that, despite our small dataset, we may have captured a central tendency in enzyme-level BNF temperature dependence. Finally, our recovery of similar BNF temperature dependence estimates across three levels of organization and terrestrial and aquatic systems suggests that the nitrogenase enzyme predictably constrains BNF rates across levels of biological organization.

Our analyses at population- and community-levels suggest that BNF temperature dependence is similar across ecological contexts and does not vary systematically with any of the ecological or evolutionary factors present in our dataset. At the population-level, this analysis was most robust, with 39 BNF-temperature responses across diverse taxonomic contexts, aquatic and terrestrial habitats, and thermally acclimated and not thermally acclimated experimental designs. Generality in the temperature dependence of BNF across aquatic and terrestrial habitats is particularly interesting, as there are major differences in the biological strategies employed by diazotrophs across these two habitats. Most notably, many diazotrophs in aquatic habitats also conduct oxygenic photosynthesis, whereas many diazotrophs in terrestrial habitats are non-photosynthetic, though they often live in close association with photosynthetic organisms. Indeed, the causes and consequences of variation in BNF rates in aquatic and terrestrial systems have historically been studied separately. Our synthesized estimate for BNF temperature dependence is quite similar to the only other synthesized estimate, which comprises only terrestrial BNF temperature responses (Houlton *et al*. 2008). Additionally, a recent global synthesis identified a similar optimum temperature for community-level BNF across aquatic and terrestrial systems, hypothesizing enzyme-level constraint as the driving mechanism (Deutsch *et al*. 2024). Our study provides empirical support for this hypothesis and contributes to a growing body of evidence that BNF in aquatic and terrestrial systems may have strong, and strikingly similar thermal constraints despite drastically different biologies.

Within metabolic scaling theory, much progress has been made in explaining effects of warming on complex systems by defining differences in the temperature dependence of key metabolic processes that are linked via resource exchange (Gilbert *et al*. 2014; Lopez-Urrutia *et al*. 2006; O’Connor *et al*. 2025). The general prediction is a thermal version of Liebig’s Law of the minimum, where a less temperature sensitive metabolism in a resource exchange network is predicted to constrain the rate of more temperature sensitive metabolisms with warming (Bozinovic *et al*. 2020; Michaletz & Garen 2024). Here, we found that across ecosystems and levels of organization, the temperature dependence of BNF rate is generally stronger than the canonical temperature dependence for photosynthesis (0.3 - 0.5 eV, Allen *et al*. 2005; Enquist *et al*. 2003; Michaletz & Garen 2024). From this, we might expect that in joint photosynthesis-BNF (C-N exchange) systems, including phytoplankton and tree-root-diazotroph symbiont systems, slower C supply from photosynthesis may limit BNF rate across a temperature gradient (Bytnerowicz *et al*. 2022; Kou-Giesbrecht *et al*. 2023). However, we find that, on average, BNF exhibits a higher temperature dependence than photosynthesis. This is despite many paired C-N exchange systems in our dataset – this factor was never an important predictor of BNF temperature-dependence in our analyses. Thus, how this thermal asymmetry is realized in C-N resource exchange systems with warming remains an open question that must be answered in order to apply our updated BNF temperature dependent estimate to global change predictions.

While the full functional response of BNF to temperature is unimodal, we quantified the thermal response in the suboptimal portion of the curve, where warming is non-stressful (Figure 1). This choice was largely practical, as there were major data constraints in our study. However, the rate of increase in BNF with warming at temperatures cooler than the thermal optimum, here temperature dependence, is also useful because it can predict how systems will respond to warming prior to crossing a thermal threshold. While other studies have emphasized the temperature at which BNF peaks (thermal optimum) as a key predictor of resource-exchange dynamics, these are primarily useful for understanding temperature thresholds where one metabolism ceases its functioning, with effects on the other metabolism (Bytnerowicz *et al*. 2022; Deutsch *et al*. 2024). Incorporating information on both sub-optimal temperature dependence and the thermal optimum is likely necessary to explain responses to warming in systems that are likely to cross the thermal optimum, including systems that exist quite near to the thermal optimum (e.g. tropical systems), or systems that experience thermal variations that surpass thermal optima. However, these approaches may not be appreciably different if environmental conditions do not exceed thermal optima or if they only exceed the thermal optimum by a small amount (Davis *et al*. 2025).

Our estimate for the temperature dependence of BNF includes data from aquatic and terrestrial habitats, symbiotic and free-living organisms, at least seven diazotroph orders (taxonomy not reported for community-level studies), and a broad geographic distribution (Figure 1). At enzyme and population levels, our dataset represents two major taxonomic groups of diazotrophs, the proteobacteria and cyanobacteria, which comprise a substantial portion of diazotrophic diversity (Koirala & Brözel 2021; Zehr *et al*. 2003). Thus, our analysis contributes a novel and robust estimate of BNF temperature dependence appropriate for use in macroecological models. However, our dataset is nonetheless geographically and taxonomically biased based on available data. System-specific models may require data from groups not represented here.

Macroecological and global models use general relationships to understand broad-scale changes across ecological contexts. Here, we present multiple lines of evidence that biological nitrogen fixation rates increase rapidly with warming in a general way across ecosystems and levels of organization, stemming from common enzyme kinetics. On average, this temperature dependence is higher than widely observed temperature dependence estimates for photosynthesis (0.3-0.5 eV). In order to clarify how this thermal asymmetry will affect carbon cycling with warming, we suggest that paired measures of photosynthetic and BNF temperature responses in controlled experiments (such as Bytnerowicz *et al*. 2022) and in natural contexts be a research priority. Further, more research is needed to determine how temperature-dependent nitrogen export processes, including denitrification, offset nitrogen inputs via BNF, and how inputs of other key nutrients including phosphorus will change with warming. This treatment of basal temperature-resource interactions as continuous and dynamic ecological processes will be central to understanding how realized warming responses in ecosystems emerge from fundamental ones in the era of global change.

## Data Availability

All data and code used to generate these results will be submitted with the manuscript in a format to facilitate double-blind review.

## Notes

### Competing Interest Statement

The authors have declared no competing interest.

### Summary of Updates

The revised manuscript focuses on enzyme-level constraint and BNF scaling as the main contribution of the manuscript.

