## Supplemental Information for "Evidence of enzyme-level thermal constraint on biological nitrogen fixation rates across systems and scales"

March 31, 2026

### Mixed effects model form

We compared hierarchical linear mixed effects using expanded versions of the general model which only included a main temperature effect (Equation 1). The full model at the population level took the following form:

$$\begin{aligned}\ln\big(B_{i,j,m}\big) = {} & \big(\hat{E}_a + \epsilon_{E_a,i,j,m}\big)\Big(\frac{1}{k\,\overline{\overline{T}}} - \frac{1}{k\,T_i}\Big) \\ & + \big(H_j\big)\Big(\frac{1}{k\,\overline{\overline{T}}} - \frac{1}{k\,T_i}\Big) + \big(O_j\big)\Big(\frac{1}{k\,\overline{\overline{T}}} - \frac{1}{k\,T_i}\Big) + \big(A_j\big)\Big(\frac{1}{k\,\overline{\overline{T}}} - \frac{1}{k\,T_i}\Big) \\ & + \Big(\ln \hat{B}_0\Big(\frac{1}{k\,\overline{\overline{T}}}\Big) + \epsilon_{B_0,i,j,m}\Big),\end{aligned} \tag{S1}$$

where where $B_{i,j,m}$ is the natural log of biological nitrogen fixation rate at temperature, $T_i$(K), for some observation, $i$, in each experimental unit, $j$, in study, $m$. Temperature is centered to the mean temperature across a level of organization, as described in the main text. $\hat{E}_a$ is the estimate of the main temperature effect across the full dataset at a level of organization and $\ln \hat{B}_0\Big(\frac{1}{k\,\overline{\overline{T}}}\Big)$ is the estimated BNF rate at the mean temperature at that level. $H_j$, $O_j$, and $A_j$ correspond to effects of habitat, Order, and thermal acclimation history on the BNF temperature response for given experimental unit. Random variation in slope and intercept attributed to experimental unit, $j$, within study, $m$, are captured in $\epsilon_{E_a,i,j,m}$ and $\epsilon_{B_0,i,j,m}$, respectively.

Table S1: List of ecological and evolutionary factors considered as possible drivers of systematic variation in temperature dependence (Ea) at each level of organization, and their definitions. Each factor was included in general temperature dependence models first as temperature interaction terms (see Equation S1). If a variable temperature dependence model was selected by AIC-based model comparison, each factor was then tested as a driver of model intercept.

| **Level of organization** | **Ecological or evolutionary factor** | **Definition** | **Levels represented** |
|---|---|---|---|
| **Enzyme** | Diazotroph species | The species that nitrogenase enzyme was isolated from | Azotobacter vinelandii, Klebsiella pneumoniae |
| **Population** | Diazotroph order | The order of diazotroph species identified in the experiment | Oscillatoriales, Pseudomonodales, Nostocales, Chroococcales, Rhodospirillales, Actinomycetales, Hyphomicrobiales |
| | Habitat | The nature of the environment where the experiment was conducted. Anything aqueous was categorized as aquatic, and non-aqueous terrestrial | Terrestrial, aquatic |
| | Oxygen accommodation strategy | Major life history strategy employed by diazotroph to shield nitrogenase from oxygen exposure, relevant primarily to phytoplankton. Non-phytoplankton were assigned ”none.” | Heterocysts, temporal separation of processes, none |
| | Symbiotic strategy | The nature of the relationship between diazotrophy and carbon acquisition. A major diagnostic was whether mass of diazotroph was measured as composite mass of diazotroph + some photosynthesizer mass (i.e. nodule biomass) in symbiotic organisms or just diazotroph mass (i.e. chlorophyll-a or total C from phytoplankton) in non-symbiotic organisms. | Symbiotic association with photosynthetic organism, no association with photosynthetic organism. |
| | Thermal acclimation history | The presence or absence of opportunity for diazotrophs at different temperature treatments to acclimate to that treatment. Exposure to an experimental treatment prior to rate measurement was categorized as sufficient for acclimation if it was expected to span several (¿ approx. 3) diazotroph generations. | Temperature acclimated, not temperature acclimated |
| **Community** | Photosynthetic association | The presence or absence of photosynthesis in close proximity to diazotrophy, whether within or across organisms. All phytoplankton, root nodules, or systems where the focal system was a phototroph were considered photosynthetically-associated. | Photosynthetically associated BNF, not photosynthetically associated BNF |

Table S2: Cross-scale model comparison of temperature effects on BNF rate. LBO is level of biological organization. Each model includes random effects on slope and intercept for experimental unit within study.

| Fixed effects structure | df | logLik | AICc | ΔAICc | Weight |
|---|---|---|---|---|---|
| temperature + LBO | 11 | -597.2 | 1217 | 0.00 | 0.82 |
| temperature * LBO | 13 | -597.0 | 1221 | 3.99 | 0.11 |
| temperature | 4 | -601.6 | 1222 | 4.82 | 0.7 |

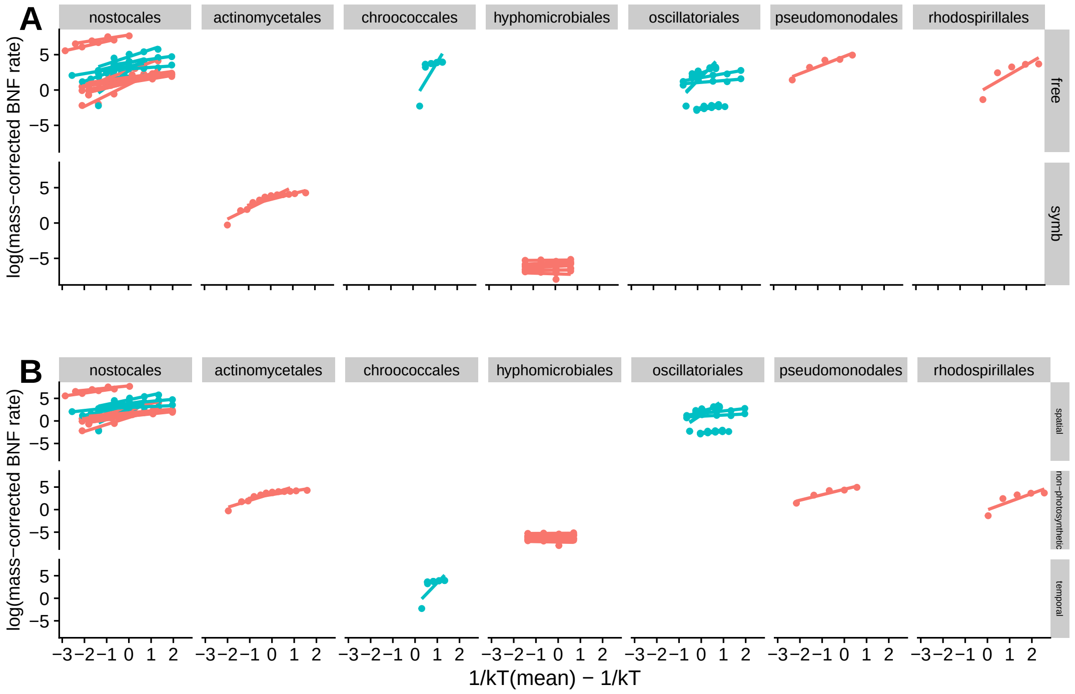


Figure S1: Distribution of population-level temperature responses across fixed effects that we were interested in testing for systematic variation in BNF temperature dependence. Each plot has columns corresponding to diazotroph order and rows corresponding to A) photosynthetic-symbiosis status (free-living or symbiotic); B) oxygen management strategy (heterocystous, non-photosynthetic, temporal). Colour corresponds to habitat type (blue = aquatic, red = terrestrial). Symbiotic status and oxygen management strategy are entirely captured by Order in our dataset.

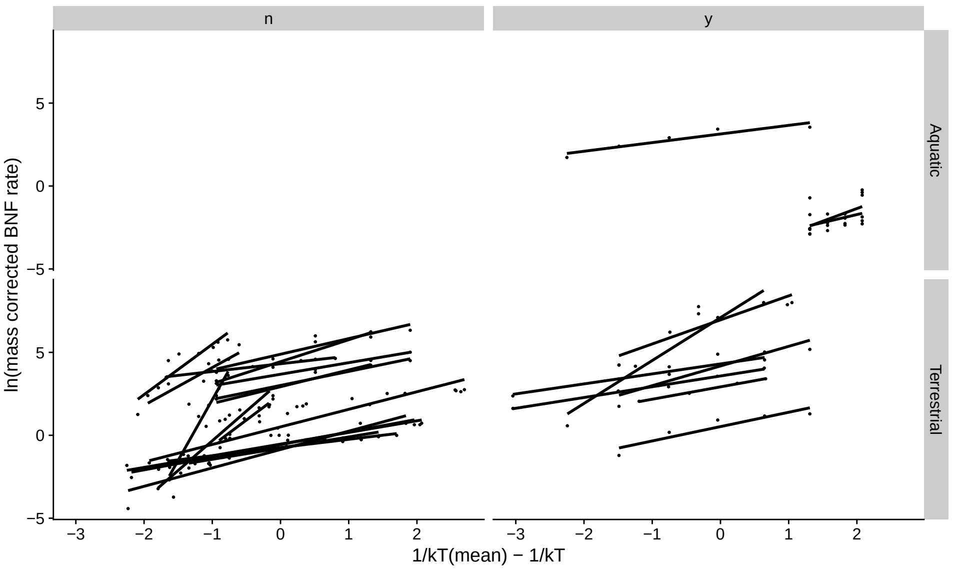


Figure S2: Distribution of community-level temperature responses across fixed effects we tested for systematic variation in BNF temperature dependence. Columns are photosynthetic association, yes ("y") or no ("n"), and rows are broad habitat type. There is insufficient data distribution across aquatic and terrestrial habitats to test for broad habitat type as a driver of BNF temperature dependence.

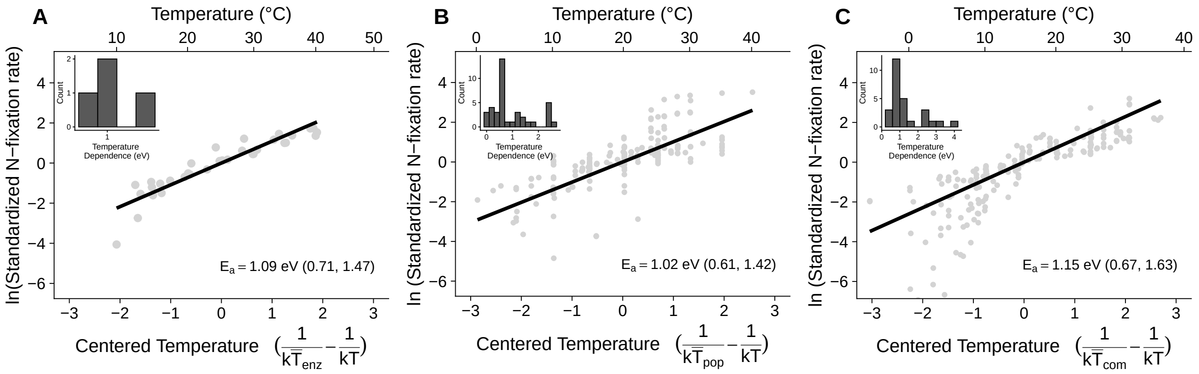


Figure S3: General temperature dependence of BNF across three levels of organization, A) enzymes, B) populations, and C) communities, based on hierarchical mixed effects model analysis. Model-predicted BNF rate-temperature relationships (black lines) and raw temperature response data (grey). Nitrogen fixation rates are standardized to the predicted rate at the mean temperature for the level of organization. Insets show the distribution of these model fits, including random effects, at each level of organization. In the enzyme-level (A) inset, the widely used BNF temperature dependence estimate ([1]) plotted as a vertical dashed line.

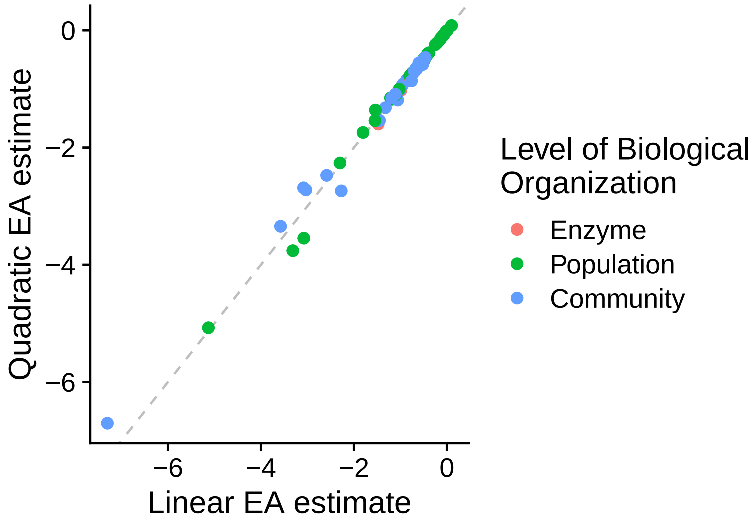


Figure S4: Relationship between temperature dependencies predicted by quadratic and linear model fits. Linear models were fit using the generalized Arrhenius model (Equation 1 in Main Text). Quadratic estimates of EA were calculated from parameters of a quadratic model $\ln\left(\frac{B}{M}\right) = q_0 + q_1(1/kT) + q_2(1/kT^2) + \epsilon$, following [2]. A linear model characterizing the relationship is overlaid on the data points.

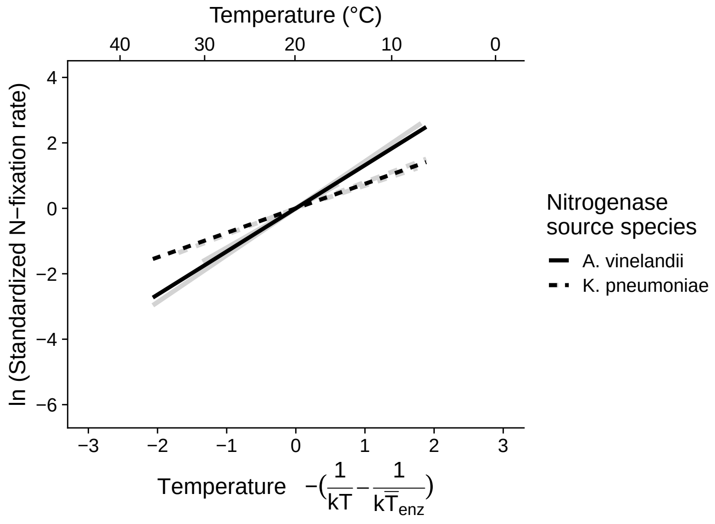


Figure S5: Predicted temperature dependence of enzyme-level BNF for *A. vinelandii* and *K. pneumoniae*, from our hierarchical mixed effects model. The whole model predicted response (black) is plotted over predictions for each individual temperature response (n = 4).
